# Geographic characterization of RPE structure and lipid changes in the PEX1-p.Gly844Asp mouse model for Zellweger spectrum disorder

**DOI:** 10.1101/2024.09.05.611330

**Authors:** Samy Omri, Catherine Argyriou, Rachel Pryce, Erminia Di Pietro, Pierre Chaurand, Nancy Braverman

## Abstract

Peroxisome Biogenesis Disorders-Zellweger Spectrum (PBD-ZSD) are a heterogenous group of autosomal recessive disorders caused by defects in *PEX* genes whose proteins are required for peroxisome assembly and function. Peroxisomes are ubiquitous organelles that play a critical role in complex lipid metabolism. Dysfunctional peroxisomes in ZSD cause multisystem effects, with progressive retinal degeneration (RD) leading to childhood blindness being one of the most frequent clinical findings. Despite progress in understanding the role of peroxisomes in normal cellular functions, much remains unknown about how their deficiency causes RD, and there is no treatment. To study RD pathophysiology in this disease, we used the knock-in PEX1-p.GlyG844Asp (G844D) mouse model of milder ZSD, which represents the common human PEX1-p.Gly843Asp allele. We previously reported diminished retinal function, functional vision, and neural retina structural defects in this model. Beyond the neural retina, structural defects in retinal pigment epithelium (RPE) have been reported in ZSD patients and murine models with single peroxisome enzyme deficiency, suggesting that RPE degeneration may contribute to overall RD progression in this disease. Here, we investigate the RPE phenotype in our PEX1-G844D mouse model, observing morphological, inflammatory, and lipid changes at 1, 3, and 6 months of age. We report that RPE cell degeneration appears at 3 months of age and worsens with time, starts in the dorsal pole, and is accompanied by subretinal inflammatory cell infiltration. We match these events with lipid remodelling using imaging mass spectrometry which allowed regional analysis specific to the RPE cell layer. We identified 47 lipid alterations that precede structural changes, 10 of which are localized to the dorsal pole. 32 of these lipid alterations persist to 3 months, with remodelling of the lipid signature at the dorsal pole. 14 new alterations occur concurrent with histological changes. Changes in peroxisome-dependent lipids detected by liquid chromatography tandem mass spectrometry (reduced docosahexanoic acid and increased very long chain lysophosphatidylcholines) are exacerbated over time. This study represents the first characterization of RPE in any animal model of ZSD, and the first *in situ* lipid analysis in any peroxisome-deficient tissue. Our findings reveal candidate lipid drivers that could be targeted to alleviate RD progression in ZSD, as well as candidate biomarkers that could be used to evaluate retinopathy progression and response to therapy.

## Introduction

Peroxisome Biogenesis Disorders-Zellweger Spectrum (PBD-ZSD) are a heterogenous group of autosomal recessive disorders due to defects in *PEX* genes whose proteins are required for peroxisome assembly and function [1]. Peroxisomes are ubiquitous organelles that play a critical role in complex lipid metabolism. Each peroxisome contains more than 50 enzymes whose functions includes β-oxidation of very long chain fatty acids (VLCFA, carbon chains ≥C22), the generation of the polyunsaturated fatty acid (PUFA) docosahexaenoic acid (DHA), and precursors for plasmalogen phospholipids [2]. Dysfunctional peroxisomes in ZSD cause multisystem effects, with progressive retinal degeneration (RD) leading to childhood blindness being one of the most frequent clinical findings. There is currently no treatment for ZSD, and management is supportive only [3].

Despite significant progress in understanding the role of peroxisomes in normal cellular functions, much remains unknown about how their deficiency causes the clinical phenotypes. Thus, we generated a knock-in (PEX1-G844D) mouse model representing the common human PEX1-p.Gly843Asp allele to study pathophysiology and develop therapies, and found that it has classic features of milder ZSD, including retinal degeneration [4]. We previously showed that this model exhibits consistently diminished cone photoreceptor function, measured from 2-32 weeks of age, with rod photoreceptor function diminishing over time, and reduced visual acuity. In addition, this model exhibits diminished cone cell numbers by 6 weeks, decreased bipolar cell numbers with age (32 weeks), and disorganization of the photoreceptor inner segment ultra structure [5, 6]. These studies are consistent with the RD observed in ZSD patients such as reduced visual acuity and cone anomalies [7].

Our previously reported ocular assessments of PEX1-G844D mice focused on retinal function and neural retina structure. However, retinal pigmentary changes are also commonly observed clinically in ZSD, suggesting the retinal pigment epithelium (RPE) is also affected in this disease [7]. Moreover, a similar RD process to that of PEX1-G844D mice was described in the *Mfp2*^*-/-*^ mouse model, which is null for a major enzyme in peroxisomal β-oxidation of very long chain fatty acids (VLCFA). This model showed earlier involvement of photoreceptor and RPE layers [8]. Selective inactivation of *Mfp2* in photoreceptors surprisingly did not result in their damage, suggesting that photoreceptor death in global *Mfp2*^*-/-*^ mice was not driven cell autonomously [9]. In contrast, selective inactivation of *Mfp2* in RPE resulted in RPE degeneration and secondary photoreceptor death [10]. Taken together, the observations in ZSD patients and *Mfp2*-deficient mouse models support the importance of peroxisome functions for RPE health. Here, we investigate the RPE phenotype in our PEX1-G844D mouse model for mild ZSD, observing morphological, inflammatory, and lipid changes over time. We report that RPE cell degeneration appears at 3 months of age and worsens with time, starts in the dorsal pole, and is accompanied by inflammatory activation. We match these events with lipid remodelling using imaging mass spectrometry (IMS) and liquid chromatography tandem mass spectrometry (LC-MSMS), identifying the earliest lipid changes that precede structural alterations, as well as lipid profiles at later disease stages.

## Materials and methods

### Mouse husbandry

G844D heterozygous mice were maintained on congenic 129/SvEv (129S6.Cg-Pex1^tm1.1Sjms^/Mmjax) and C57BL/6N (B6.Cg-^Pex1tm1.1Sjms^/Mmjax) backgrounds. As >80% of PEX1-G844D homozygotes uniformly die before weaning on either congenic background [11, 12] all experiments were performed on mice with a stable mixed background of 50% C57Bl/6 and 50% 129Sv/Ev. The C57Bl/6 strain used is negative for the Rd8 mutation of the *Crb1* gene. The Rd8 mutation was removed by crossing with the Crb1^cor^ C57BL/6N strain (JAX 022521). To minimize genetic variation, we used the F1 generation of two congenic PEX1-G844D heterozygote parents (♂C57Bl/6 x ♀129Sv/Ev, or ♀C57Bl/6 x ♂129Sv/Ev). We observed no effects of parental imprinting or sex. Nearly all (>90%) PEX1-G844D homozygous F1 progeny survive past weaning, after which we observe no effect of the mutation on life expectancy. Mice were housed at the RI-MUHC Glen site animal care facility with *ad libitum* access to food and water. All experiments were performed at the RI-MUHC Glen site and approved by the Research Institute of the McGill University Health Centre Animal Care Committee. Euthanasia was performed using CO2 under isoflurane anaesthesia (5% isoflurane in oxygen until loss of consciousness, immediately followed by CO2 at maximum flow rate, 4 LPM). Both males and females were used for all experiments, and wild-type and PEX1-G844D heterozygous mice used as littermate controls. There were no phenotypic differences based on sex or control genotype. Genotyping was performed as previously described [5].

### Retinal flatmount preparation and immunofluorescence

Mouse eyes were enucleated and fixed in 4% formalin for 5 min at room temperature, at which point a hole was made in the cornea with a needle (27G), and eyes fixed for additional 25 min. Fixed eyes were sectioned at the limbus; the anterior segments were discarded. The posterior eye cups consisting of the neural retina/RPE/choroid/sclera complex were collected and the neural retina was carefully detached from RPE/choroid/sclera to be prepared separately for experiments.

<u>RPE/choroid/sclera flatmounts</u> were treated with PBS solution containing 0.1% Triton X-100 for45 min. Specimens were incubated overnight at 4°C with 1:400 TRITC phalloidin (ECM Biosciences, PF7551 Versailles, KY) for RPE actin cytoskeleton labelling. Primary antibody used was 1:400 rabbit anti-human IBA1 (Wako 019-19741, Richmond, VA) with secondary antibody 1:450 anti-rabbit Alexa Fluor 488 (Invitrogen A21206, Waltham, MA). Immunofluorescence was analyzed and images acquired using a Zeiss LSM780 laser scanning confocal microscope. We counted the number of RPE cells or IBA1 positive cells in 8 fields of view spanning the four poles of each RPE flatmount (excluding the centre, containing the optic nerve head) to quantify RPE degeneration or inflammation, respectively, and reported this as cells/mm^2^.

<u>Neural retina flatmounts</u> were treated with PBS solution containing 0.1% Triton X-100 and 10% FBS (saturation buffer) for 45 min. Cone photoreceptors were stained overnight at 4°C with 1:500 fluorescein peanut agglutinin (Vector Laboratories, Burlingame, CA) diluted in saturation buffer. Imaging was performed as above.

### Retinal immunohistochemistry

Eye cups from PBS-perfused mice were fixed 3 hr. in 10% formaldehyde, incubated in 10% (30 min. on ice), 20% (1 hr. on ice), and 30% (4°C overnight) sucrose in 0.1M PB, then embedded and frozen in frozen section compound (VWR, Mississauga, Canada). 5µm retinal cryo-sections were blocked (1% normal goat serum, 0.1% Triton X-100, 10% bovine serum albumin (BSA) in PBS) for 1 hr, washed, incubated at 4°C overnight with 1:400 rabbit anti-human GFAP (Abcam ab7260 Waltham, MA) in incubation buffer (0.1% Triton X-100, 10% BSA in PBS). Following primary antibody incubation, sections were washed and incubated 90 min. with 1:450 anti-rabbit Alexa Fluor 594 (Invitrogen A21207, Waltham, MA) and washed again. Coverslips were mounted using ProLong Gold antifade reagent with DAPI (Invitrogen, Burlington, CA) and retinas were visualized by confocal microscopy as above.

### Matrix-assisted laser desorption ionization (MALDI) mass spectrometry

RPE was harvested from sets of three PEX1-G884D and WT mice at 30 and 90 days of age. The tissues were flat-mounted onto conductive indium tin oxide-coated microscope slides (Delta-Technologies Ltd. Loveland, CO, USA).

#### Silver-assisted laser desorption ionization (LDI) mass spectrometry imaging (MSI)

Silver-assisted LDI MSI was performed after silver sputter deposition for 30s at 80mA and 0.02 mbar of argon partial pressure on the RPE sections using a Cressington 308R sputter coater system (Ted Pella Inc. Redding, CA, USA) as previously described [13]. Sections were then analyzed in positive ion mode with reflectron geometry using ultrafleXtreme MALDI-TOF mass spectrometer (Bruker Daltonics. Billerica, MA, USA). Source parameters including delayed extraction and laser energy were optimized for maximum mass resolving power and signal to noise. Automated MSI data acquisition was performed at a spatial resolution of 150μm averaging 300 shots per coordinates. MSI data acquisition was performed using the Bruker Daltonics flexImaging V4.1software.

#### Dual-polarity MALDI MSI

Dual-polarity MALDI MSI was performed after 1,5-diaminonapthlene matrix sublimation using a home-built system onto the RPE flatmounts for 7min at 140°C as previously described [14]. The tissues were then analyzed by MSI in reflectron mode under optimized ion source settings using the ultrafleXtreme system at a 150μm spatial resolution averaging 150 laser shots in positive and 350 laser shots in negative ion per coordinate. The tissues were first imaged in positive ion mode. For negative ion mode MSI, the coordinate acquisition grid was offset by 75μm in both the x and y directions allowing resampling of the same sections.

#### MSI data processing

MSI data processing was performed with the Cardinal V3 software within the R V4.2.3, environment along with some lab generated and base R functions [15]. Data was normalized by the total ion current and binned with a tolerance of 0.2Da. For each considered m/z signal, average intensity values were generated per section or per RPE areas (dorsal and ventral). To reduce the effect of outlier pixels, the top and bottom 10% of the MSI data was not considered in the determination of mean values for each region.

#### Tandem mass spectrometry

MS/MS data acquisition for fatty acids and lipids was performed on a TimsTOF Flex mass spectrometer (Bruker Daltonics. Billerica, MA, USA). Identifications only based on exact mass only (mass accuracy <2ppm) were also determined using this instrument.

### LC-MSMS analysis

For liquid chromatography-tandem mass spectrometry (LC-MS/MS) analysis, RPE or neuroretina was homogenized in PBS using a mini pestle. 2:1 chloroform/methanol containing 0.05% butylhydroxytoluene (BHT) was added to homogenized tissue in a glass tube and incubated on an orbital shaker at room temperature for 2 hrs. Samples were centrifuged at 2500rpm for 10 min. and the supernatant was transferred to a clean glass tube. The supernatant was washed with 0.2 volumes of purified water, mixed and centrifuged at 2000 rpm room temperature for 5 min. to separate the two phases. The upper phase was removed and the lower phase washed with folch theoretical upper phase. (3:48:47 chloroform:methanol:water). Samples were mixed and centrifuged at 2000 rpm for 5 min. The upper phase was removed and the lower phase dried under nitrogen. The dried lipid was dissolved in 3:2 hexane:isopropanol containing 10 ng of each internal standard, 16:0-D4 lyso-PAF (20.6 pmol) and D4-26:0-lyso-PC (15.6 pmol). Samples were filtered by centrifugation through Costar spin-X centrifuge tube filters (Corning, NY) for 5 min. Filtrates were analyzed in Verex auto-sampler vials (Phenomenex, Torrance, CA). A 2.1 × 50 mm,1.7 µm chromatography column and a Waters (Milford, MA) TQD (Triple Quadrupole Mass Spectrometer) interfaced with an Acquity UPLC (ultra-performance liquid chromatography) was used in positive ion electrospray (ESI)-MS/MS ionization. Solvent systems were: mobile phase A= 54.5% water/45% acetonitrile/0.5% formic acid, mobile phase B = 99.5% acetonitrile/0.5% formic acid with both solutions containing 2 mM ammonium formate. Initial solvent conditions were 85% mobile phase A/15% mobile phase B followed by a gradient from 15% to 100% mobile phase B over a period of 2.5 min., held at 100% mobile phase B for 1.5 min. before reconditioning the column back to 85% mobile phase A/ 15% mobile phase B for 1 min. at a solvent rate of 0.7 ml/min. A column temperature of 35°C and an injection volume of 5ul was used for analysis. Ethanolamine plasmalogens were detected by multiple reaction monitoring (MRM) transitions representing fragmentation of [M+H]+ species to m/z 311, 339, 361, 385, 389 for compounds with 16:1, 18:1, 20:4. 22:6 and 22:4 at the sn-2 position, respectively. Lysophosphatidylcholine (LysoPC) species were detected by multiple reaction monitoring (MRM) transitions representing fragmentation of [M+H]+ species to m/z 104. Reagents used were authentic plasmalogen standards, tetradeuterated internal standards 26:0-D4 lysoPC (Avanti Polar Lipids, Alabaster, Alabama), 16:0-D4 lyso PAF (Cayman Chemical Company, Ann Arbor, MI) and Optima grade solvents (methanol, acetonitrile, chloroform, water) (Fisher Scientific, Waltham, MA), formic acid (Honeywell Fluka), ammonium formate (Sigma-Aldrich), and PBS (Thermo Fisher Scientific, Waltham, MA).

### Statistical Analysis

Results are expressed as mean ± SD (Standard Deviation). Statistical significance was calculated with unpaired Student’s t-test to compare 2 groups, assuming normal distribution and homogeneity of variances. Comparisons between more than 2 groups were performed on normally distributed data using one-way ANOVA and Tukey’s multiple comparisons test. Statistical significance was set based on P value: *P < 0.05, **P < 0.01, ***P < 0.001, ****P < 0.0001. All experiments were repeated at least 3 times. Statistical analysis was performed using GraphPad Prism 10.0 software

## Results

### Histological analysis of PEX1-G844D RPE and identification of early lipid changes

To investigate the initial stages of retinal changes in PEX1-G844D mice, we performed histological analyses in combination with imaging mass spectrometry (IMS) of the retinal pigment epithelium (RPE) at 1 month-old.

#### Retinal pigment epithelium (RPE) morphology

We examined cell morphology in RPE flatmounts by labeling polymerized actin filaments (F-actin) with phalloidin-TRITC to visualize the cortical actin cytoskeleton delineating the RPE cells. Confocal imaging analysis revealed the characteristic “honeycomb” shape of the RPE in PEX1-G844D mice, with no apparent difference compared to littermate controls (**Figure 1A**). We confirmed this by scoring the number of RPE cells in 8 fields of view spanning the four poles of each RPE flatmount (excluding the optic nerve area) and reported this as cells per surface area. Quantification confirmed normal integrity of the PEX1-G844D RPE layer with no significant lesions compared to age-matched control mice (**Figure 1B**).

**Figure 1.**
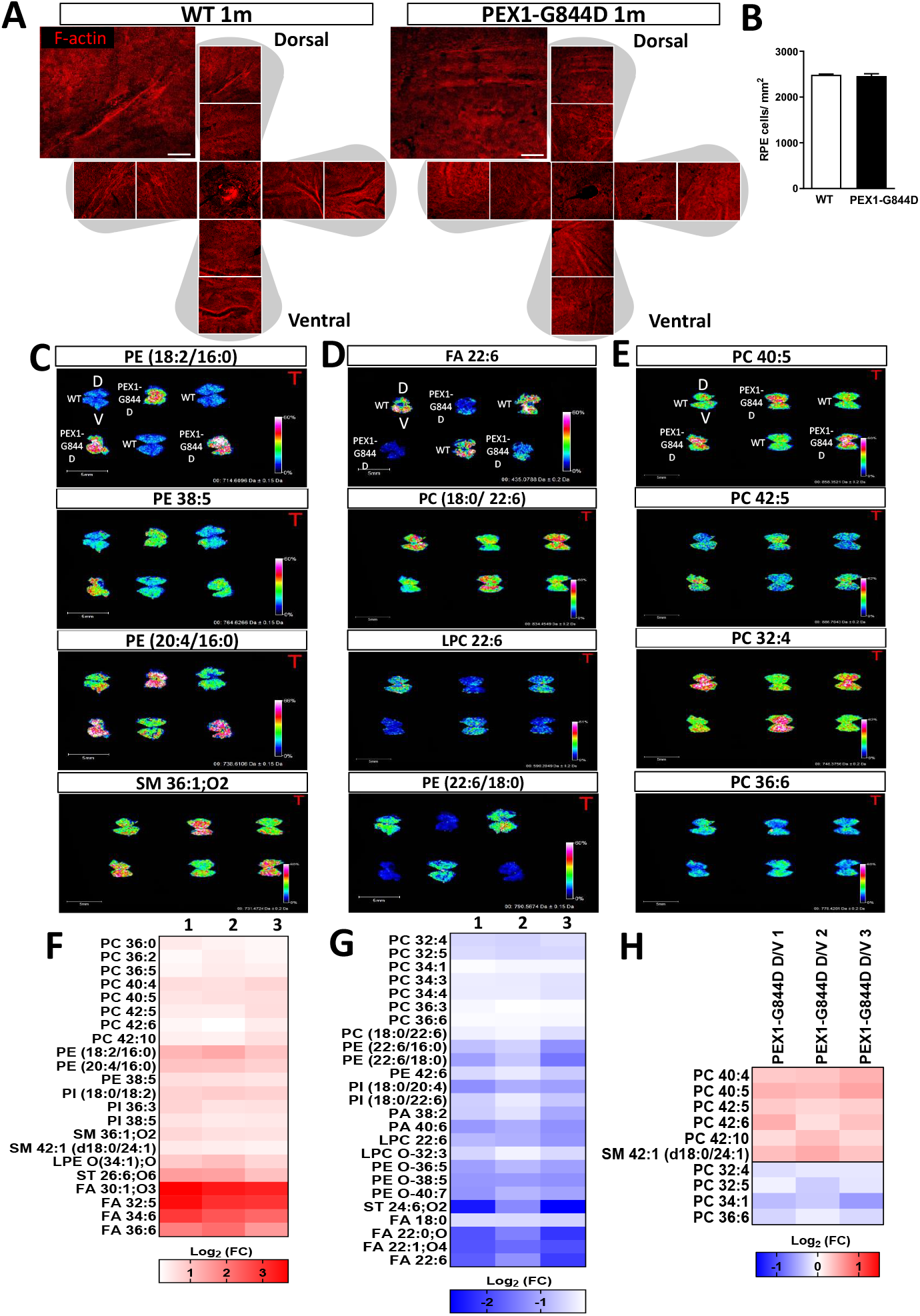
PEX1-G844D induces early lipid changes in RPE at 1 month of age. **A:** Confocal imaging of RPE flatmounts from 1-month-old WT and PEX1-G844D mice counterstained with TRITC-phalloidin (F-actin marker). The four-petal flower (grey) represents RPE tissue prepared for flat mounting. Nine confocal microscope images arranged in dorso-ventral orientation were taken to visualize tissue integrity. A zoomed image is included at the top right of each RPE to facilitate observation of cell morphology. Scale bar: 100 μm. **B:** Quantification of RPE cell number from WT and PEX1-G844D RPE flatmounts (N=3 per group). **C-D:** Examples of IMS analysis (6 RPE flat mounts from 1 month-old mice: 3 WT in V position and 3 PEX1-G844D in inverted V position) showing 12 lipids identified with **(C)** increased density, **(D)** reduced density and **(E)** increased density specifically in the dorsal pole of the RPE. **F-H:** Heatmap of selected lipid species with **(F)** significant increase, **(G)** or significant decrease compared to control. **(H)** Significant lipid changes in dorso-ventral ratio in RPE flatmounts of PEX1-G844D. Each column (1, 2 and 3) represent a ratio of IMS analysis of one PEX1-G844D RPE sample/ average of IMS analysis of WT RPE. Each row corresponds to a lipid species. **(F)** Heatmap of lipids with significant increase compared to control and showing significant change in dorso-ventral ratio in RPE flatmounts of PEX1-G844D. The heat map color scale indicates the highest lipid density in dark red and the lowest lipid density in dark blue.

#### Early lipid changes in PEX1-G844D RPE tissue at 1 month

In the absence of structural alterations to the RPE, we next examined the tissue for alterations in lipid composition. Identifying early RPE lipid changes would inform the pathophysiology of PEX1-G844D-induced photoreceptor dysfunction, given the importance of RPE membrane trafficking for photoreceptor support [16]. Changes in membrane lipid composition can impact membrane fluidity and the function of membrane-bound proteins, which are crucial for nutrient transport, waste removal, and the visual cycle. Disruptions to lipid metabolism due to peroxisome deficiency in the RPE could thus lead to photoreceptor dysfunction and contribute to retinal disease.

To identify PEX1-G844D-induced lipid changes in the RPE at 1 month of age, we used imaging mass spectrometry (IMS), to map the distribution of multiple lipid species in RPE tissue. This technique enables lipid detection exclusively in the RPE, separate from the choroid and sclera that would be present in RPE tissues dissected for traditional liquid chromatography tandem mass spectrometry (LC-MSMS) analysis. Lipid profiling identified more than seventy lipids among phosphatidylcholines (PC), phosphatidylethanolamines (PE), phosphatidylinositol (PI), phosphatidyl acids (PA), and sphingomyelins (SM) (**Supplemental tables S1 and S2**).

Comparison of IMS data from PEX1-G844D and WT RPE flatmounts revealed clear differences in relative densities in over forty lipids (**Figures 1C-E**). Lipid species with a higher density distribution pattern in PEX1-G844D RPE compared to control are shown in **Figure 1C**, and lower density in **Figure 1D**. These changes suggest that RPE membrane composition is altered in PEX1-G844D mice compared to wild-type controls, even at an early age and in the absence of structural defects. Unexpectedly, certain lipid species showed differential distribution within the PEX1-G844D RPE (**Figure 1E**). Among them, PC 40:5 and PC 42:5 were increased, whereas PC 32:4 and PC 36:6 were decreased, exclusively in the dorsal pole. This phenomenon did not occur in control RPE flatmounts, where lipid distribution was homogeneous throughout the tissue.

All the lipid changes were quantified (**Supplemental Table S1**) and analysis revealed 16 lipids, of which three [PC 34:1, PE (22:6/18:0), PI (18:0/20:4)] had lower density and four [PC 36:0, PC 36:2, PC 36:5, PE (20:4/16:0)] had higher density in PEX1-G844D RPE compared to WT RPE. Heat maps of whole RPE tissues are presented for all these lipid species that were statistically higher (**Figure 1F**) or lower density (**Figure 1G**) in PEX1-G844D *versus* controls.

As expected with peroxisome dysfunction, IMS analysis revealed a significant increase in very long-chain fatty acids (VLCFAs) including: FA 30:1;O3, FA 32:5, FA 34:6 and FA 36:6. We also observed a significant decrease in plasmalogens, such as LPC O-32:3, PE 16:0p20:4, PE O-38:5, PE O-40:7 and PA O-42:1. In addition, we detected a reduction in docosahexaenoic acid (C22:6, DHA) and multiple lipids with DHA incorporated, notably LPC 22:6, LPA 22:6, PC (18:0/ 22:6), PE (16:0/ 22:6), PE (18:0/ 22:6), PI (18:0/ 22:6) and PG (22:6/ 22:6).

Based on the accumulation of certain lipids observed with IMS in the dorsal pole of the PEX1-G844D compared to control RPE (**Figure 1E**), we analyzed whether this different distribution between the poles was statistically significant. We found a significant accumulation of PC 40:4, PC 40:5, PC 42:5, PC 42:6 and SM 42:1 in the RPE membrane in the dorsal compared to ventral pole (**Figure 1H**). In contrast, there was a significant decrease in PC 32:4, PC 32:5, PC 34:1 and PC 36:6 in the dorsal pole (**Figure 1 H**).

### Lipid dysregulation induced by the PEX1-G844D mutation precedes retinopathy

Knowledge of early lipid dysregulation in the PEX1-G844D mouse RPE prompted further investigation into the progression of RPE changes in this model. We expanded our histological study of RPE flatmounts to include tissues from 3-month-old mice. Staining of cortical actin filaments revealed profound structural changes in PEX1-G844D mice at this age (**Figure 2**). Compared to WT control (**Figure 2A**), PEX1-G844D RPE tissue exhibited profoundly enlarged cells with disorganized shape (**Figure 2B**). This is a hallmark of RPE degeneration, wherein cell death results in enlargement of remaining cells to preserve the external blood retinal barrier function [17]. Most strikingly, these structural abnormalities occurred mainly in the dorsal pole, where certain lipids specifically accumulated at 1 month of age. Cell quantification showed a 35% reduction in the number of RPE cells per overall RPE surface area in PEX1-G844D compared to controls (**Figure 2C**, left). In the dorsal area specifically, RPE cell density was reduced by 63% in PEX1-G844D mutants, while there was no difference between mutants and controls in the ventral area (**Figure 2C**, right). In addition, we observed the presence of blebs on PEX1-G844D RPE plasma membranes (**Figure 2B**, middle), indicative of RPE cellular stress [18].

**Figure 2.**
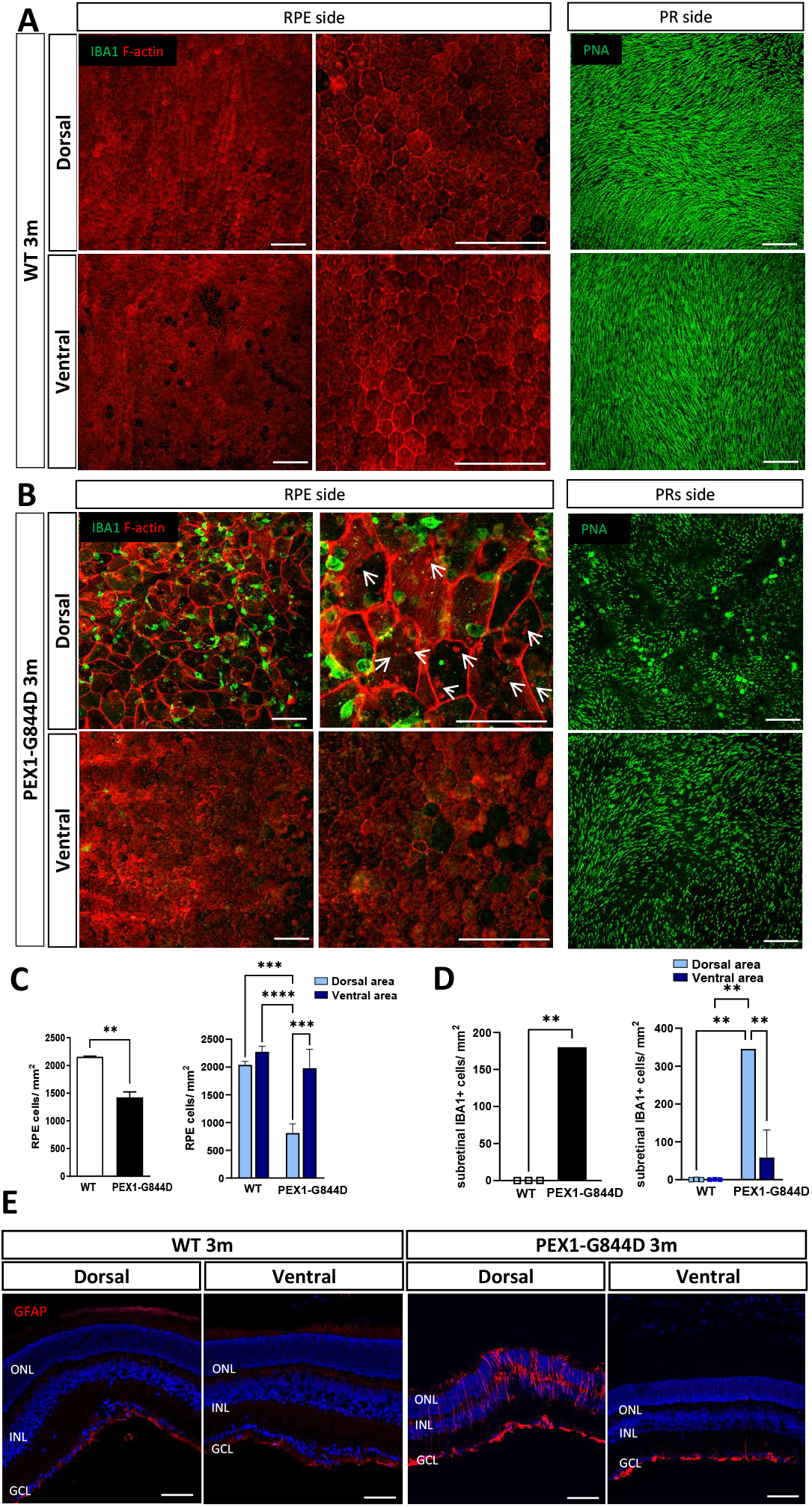
Early PEX1-G844D retinopathy phenotype at 3 months. **A-B:** Confocal imaging of dorsal and ventral areas of 3 month-old **(A)** WT littermate control **(B)** and PEX1-G844D RPE flatmounts immunostained with IBA1 antibody, a microglia/ macrophage marker (green) and counterstained with TRITC-phalloidin (red) (left and middle panels). White arrows show the formation of blebs within RPE cells (middle panel). Neuroretina flatmounts oriented with photoreceptors face up (PR side) were counterstained with peanut agglutinin (PNA), a cone marker (green) (right panels). **C:** Quantification of RPE cell number per mm^2^ in whole RPE tissue (left graph) and in dorso-ventral areas (right graph). (N=3, **p< 0.01, ***p< 0.001, ****p< 0.0001). **D:** Quantification of subretinal IBA1 positive cells per mm^2^ on the whole RPE (left graph) and in dorso-ventral areas (right graph) (N=3,**p< 0.01). **E:** Confocal imaging of dorsal and ventral poles of retinal cryosections from 3 month-old WT and PEX1-G844D mice, immunostained with GFAP antibody (red) counterstained with DAPI (blue). ONL: outer retinal layer; INL: inner nuclear layer; GCL: ganglion cell layer. Scale bar: 100 μm.

Subretinal inflammation is a strong component of retinopathy progression in different RD models (age-related macular degeneration (AMD), diabetic retinopathy (DR)) [19-23], including other models of RD associated with lipid dysregulations such as the *Mfp2*^*-/-*^ and *Mfsd2a*^*-/-*^ mouse models [8, 24]. Thus, we investigated whether the structural changes in PEX1-G844D RPE are associated with subretinal inflammation. Moreover, the presence or absence of inflammation and its impact on photoreceptors would allow us to better understand and characterize the development of retinal disease in our model. Immunofluorescent microscopy of RPE flatmounts incubated with a marker of microglia/macrophages (IBA-1) showed increased infiltration of inflammatory cells in the PEX1-G844D subretinal space, indicating an active inflammatory response (**Figure 2A,B**). On average, there were 175 IBA-1+ cells per mm^2^ in the PEX1-G844D RPE, and complete absence of these inflammatory cells in control tissues (**Figure 2D**, left). Quantification according to the geographic area showed an accumulation of IBA1+ cells mainly in the dorsal pole (300 cells/mm^2^) compared to the ventral pole (50 cells/mm^2^) of the retinal tissue (**Figure 2D**, right). In parallel, histological examination of the photoreceptor side of neuroretina flatmounts stained with a marker of cone extracellular matrix (peanut agglutinin, PNA) revealed decreased cone outer segment density. This occurred most prominently at the dorsal pole, in the same area as, and facing, the damaged RPE (**Figure 2A,B**). Furthermore, immunofluorescent analyses of retinal cryosections showed increased levels of glial fibrillary acidic protein (GFAP), an indicator of gliosis, only in the dorsal pole (**Figure 2E**), confirming an acute retinal stress condition in this specific zone of the PEX1-G844D retina.

### Onset of retinal alterations and correlation with lipid dysregulation

Our findings suggest that lipid dysregulation at earlier stages may contribute to the observed inflammation and retinal damage at 3 months of age in PEX1-G844D mice. In this context, we proceeded to analyze the evolution of distribution for the lipids identified previously at 1 month of age using IMS, within this newly characterized inflammatory environment.

Lipid profiling in 3-month-old RPE using IMS showed further changes in lipid composition, quantified and referenced in **Supplemental table S2**. More than 40 lipid species showed statistically significant changes in PEX1-G844D RPE compared to control RPE (**Figure 3A**). Among the twenty most abundant lipid species detected by IMS, 6 showed increased density in mutants: PC 34:2, PC 36:4, PC 36:5, PC (16:1/22:6), PC 40:7 and PC 40:8, whereas five showed decreased density: PC 34:0, PC 34:1, PC 36:1, PI (18:0/20:4) and SM 34:1; O2, suggesting a trend towards increasing polyunsaturated fatty acids. *In situ* examples of this differential lipid expression in PEX1-G844D and control RPE are illustrated in **Figure 3B**, with increased lipid species presented in the left column, and decreased lipid species in the right column.

**Figure 3.**
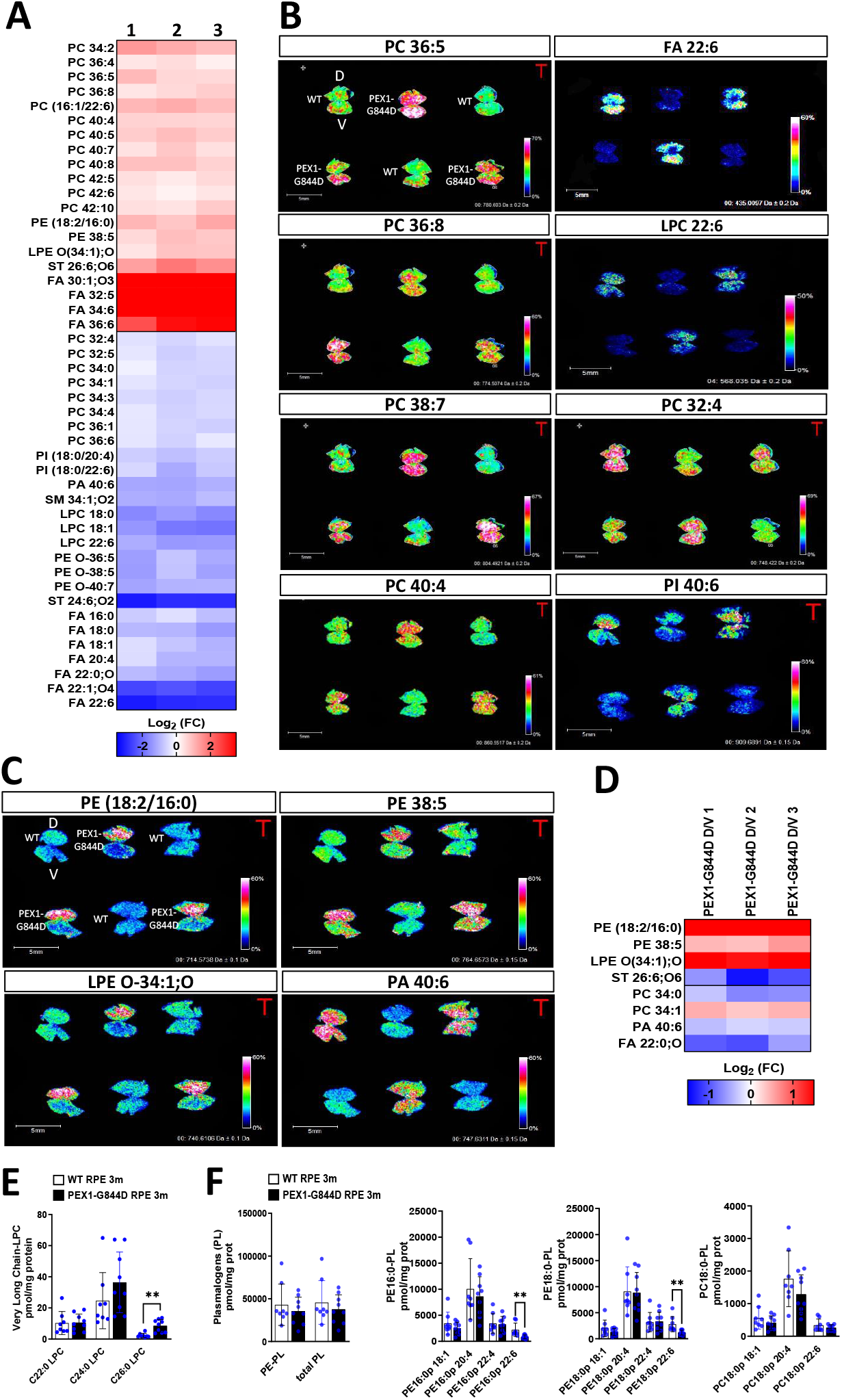
RPE lipid signature associated with neuroinflammation in PEX1-G844D at 3 months of age. **A:** Heatmap of selected lipid species with significant changes (p< 0,05) compared to control. Each column (1, 2 and 3) represents a ratio of IMS analysis of one PEX1-G844D RPE sample/ average of IMS analysis of WT RPE. Each row corresponds to a lipid species. **B:** IMS analysis (6 RPE flatmounts from 3 month-old mice: 3 WT in V position and 3 PEX1-G844D in inverted V position) showing 4 lipids identified with increased density (left column) or 4 other lipids with reduced density (right column) . **C:** IMS analysis showing increased density specifically in the dorsal pole of the RPE. **D:** Heatmap of lipids with significant increase compared to control and showing significant change in dorso-ventral ratio in RPE flatmounts of 3 month-old PEX1-G844D mice. The heat map color scales indicate highest lipid density in dark red and lowest lipid density in dark blue. **E, F:** Histograms presenting the levels of **(E)** Very Long Chain-Lysophosphatidylcholines (VLC-LPCs) and **(F)** phosphatidylethanolamine (PE), phosphatidylcholine (PC) plasmalogens measured by LC/MSMS in RPE tissue from WT and PEX1-G844D mice. (N=8, **p<0,01).

Most of the specific lipid species changes observed at 1 month persisted to 3 months, including the decrease of DHA and DHA-associated lipids (LPC 22:6, PI (18:0/22:6)), and an exacerbated increase in VLCFAs (FA 30:1; O3, FA 32:5, FA 34:6 and FA 36:6). These species were homogeneously altered throughout the RPE tissue. These results indicated a sustained increase of polyunsaturated lipid species in PEX1-G844D RPE. Three of the lipid species increased evenly throughout the RPE at 1 month, but at 3 months showed a higher accumulation in the dorsal pole: PE (18:2/16:0), PE 38:5 and LPE O-34:1; O (**Figure 3C,D**). These lipids, which increase at the site of histological changes at 3 months, could be markers of or involved in active tissue damage and inflammation.

Interestingly, the lipids previously accumulated in the dorsal pole at 1 month were still increased compared to control at 3 months (excepted for SM 42:1), but spread homogeneously throughout the whole RPE tissue. Early accumulation in the dorsal pole followed by a more generalized increase indicates that these lipids may accompany or follow the spread of tissue damage and inflammation.

Our IMS data revealed significant alterations in the lipid composition of the RPE membrane, associated with the development of RPE inflammation and lesions. We aimed to determine whether these findings could be associated with the presence of lipid biomarkers characteristic of ZSD commonly detected by LC-MSMS. Our LC-MSMS analysis of very long chain LPCs and PE-, PC-plasmalogens (PE-PLs, PC-PLs) in the isolated RPE-choroid-sclera complex revealed a 3-fold increase in C26:0 LPC levels in PEX1-G844D tissues (**Figure 3E**), while overall (total) plasmalogen levels were not significantly altered. In contrast, a specific analysis of DHA-containing plasmalogens (C22:6) showed a 50% decrease of PE 16:0p22:6 and PE 18:0p22:6 (**Figure 3F**). This observation corroborates the IMS analyses, which showed a consistent decrease in DHA-associated lipids. This complementary approach confirmed the simultaneous presence of ZSD biomarkers with the lipid changes revealed by IMS and the retinopathy phenotype.

### PEX1-G844D retinopathy extends to the whole RPE after 6 months

To understand the long-term progression of inflammation and tissue damage in the PEX1-G844D mouse model, we extended histological analysis of the RPE and neural retina to 6 months. In addition, these studies would show whether the lipid changes that migrated from the dorsal to ventral pole at 3 months also preceded inflammation and tissue damage. Indeed, RPE flatmounts at 6 months of age demonstrated structural changes with enlarged and disorganized cells accompanied by subretinal inflammation throughout the entire RPE tissue (F-actin and IBA-1 staining, **Figure 4A**). Quantification of these changes showed a 60% reduction of RPE cells in PEX1-G844D compared to control mice, and an average of 180 IBA-1+ cells per mm^2^ (**Figure 4B**). This is consistent with spatial lipid changes preceding structural events. Matching the widespread RPE degeneration, cone outer segment density (visualized using PNA) declined across the neural retina (**Figure 4C**). Pan-retina photoreceptor cell loss occurred by this stage, evidenced by a thinning of the outer nuclear layer (ONL), although this was far more pronounced in the dorsal pole (**Figure 4D**). At 6 months, gliosis (GFAP) was present in both dorsal and ventral poles (**Figure 4D**). Changes in peroxisome biochemical metabolites in the PEX1-G844D RPE-choroid-sclera complex were exacerbated from 3 to 6 months; LC-MSMS analysis showed a 10-fold increase in C26:0 LPC levels in the RPE-choroid-sclera complex (**Figure 4E**), and a 65% decrease in the DHA-containing plasmalogens PE 16:0p22:6 and PE 18:0p22:6 (**Figure 4F**).

**Figure 4:**
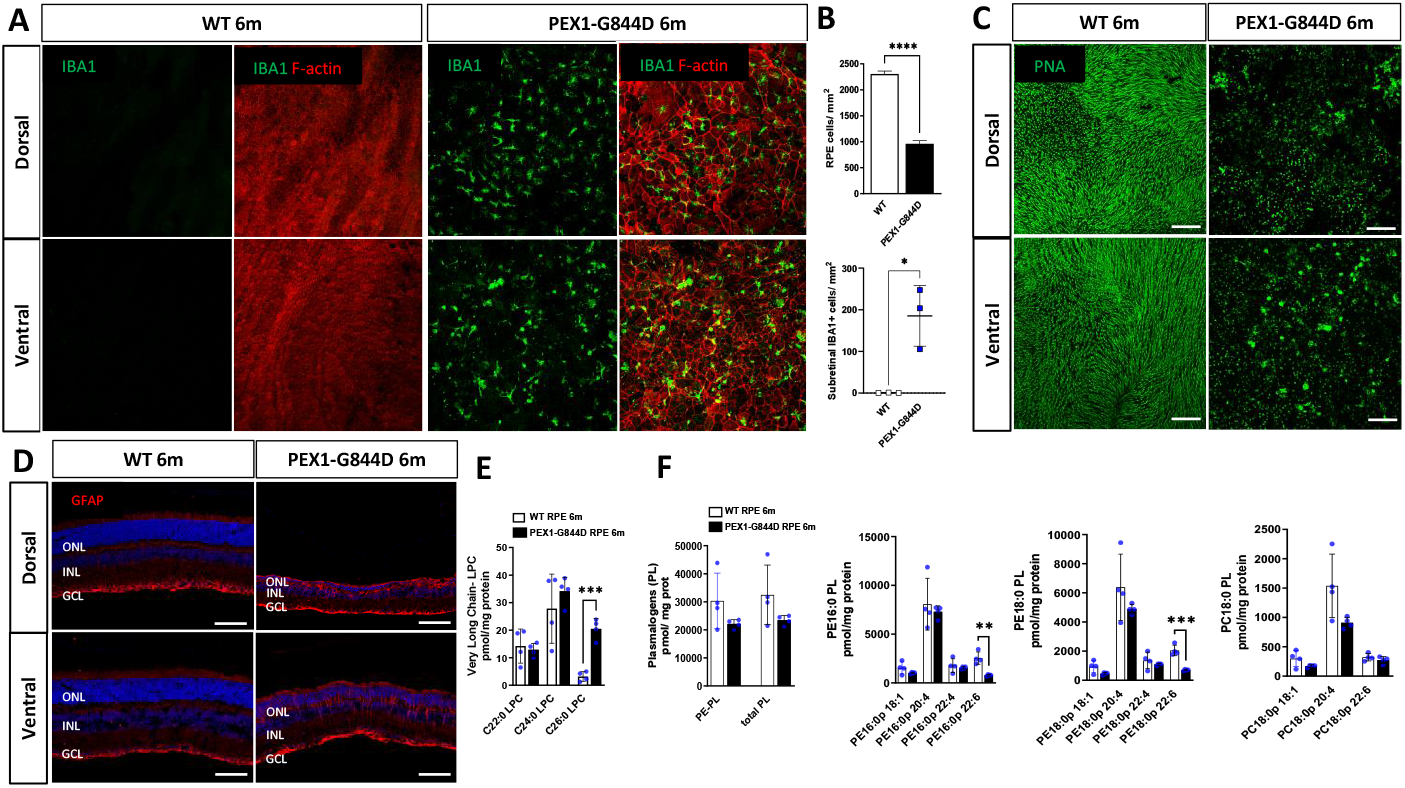
Later stage PEX1-G844D retinopathy phenotype at 6 months. **(A)** Confocal imaging of dorsal and ventral areas of 6 month-old WT and PEX1-G844D RPE flatmounts immunostained with IBA1 antibody, a microglia/ macrophage marker (green) and counterstained with TRITC-phalloidin (red). **(B)** Quantification of RPE cell number per mm^2^ in whole RPE tissue (top graph) and quantification of subretinal IBA1 positive cells per mm^2^ on the whole RPE (bottom graph) (N=3,*p< 0.05, ****p< 0.0001). **(C)** neuroretina flatmounts oriented with photoreceptors face up counterstained with peanut agglutinin (PNA), a cone marker (green). **(D)** Confocal imaging of dorsal and ventral poles of retinal cryosections from 6 month-old WT and PEX1-G844D mice, immunostained with GFAP antibody (red) counterstained with DAPI (blue). ONL: outer retinal layer; INL: inner nuclear layer; GCL: ganglion cell layer. Scale bar: 100 um. **E, F:** Histograms presenting the levels of **(E)** VLC-LPCs and **(F)** phosphatidylethanolamine (PE), phosphatidylcholine (PC) plasmalogens measured by LC/MSMS in RPE tissue from WT and PEX1-G844D mice. (N=8, **p<0.01, ***p<0.001).

### Comparison of PEX1-G844D-induced-lipid change with other RD mouse models with lipid dysregulation

We illustrated all the characteristic lipid signatures of the PEX1-G844D RPE in a Venn diagram (**Figure 5A**). We found that 47 lipids were precociously affected during the first month. 15 of them were found to be modified only at early stages of the disease, before the onset of retinal damage. The intersections of the diagram show that 32 lipids were significantly modified and persisted over the first 3 months in PEX1-G844D RPE. These include the lipid species specifically accumulated in the dorsal pole at 1 and 3 months. An additional 14 lipids showed significant changes only once the retinopathy phenotype appeared.

**Figure 5.**
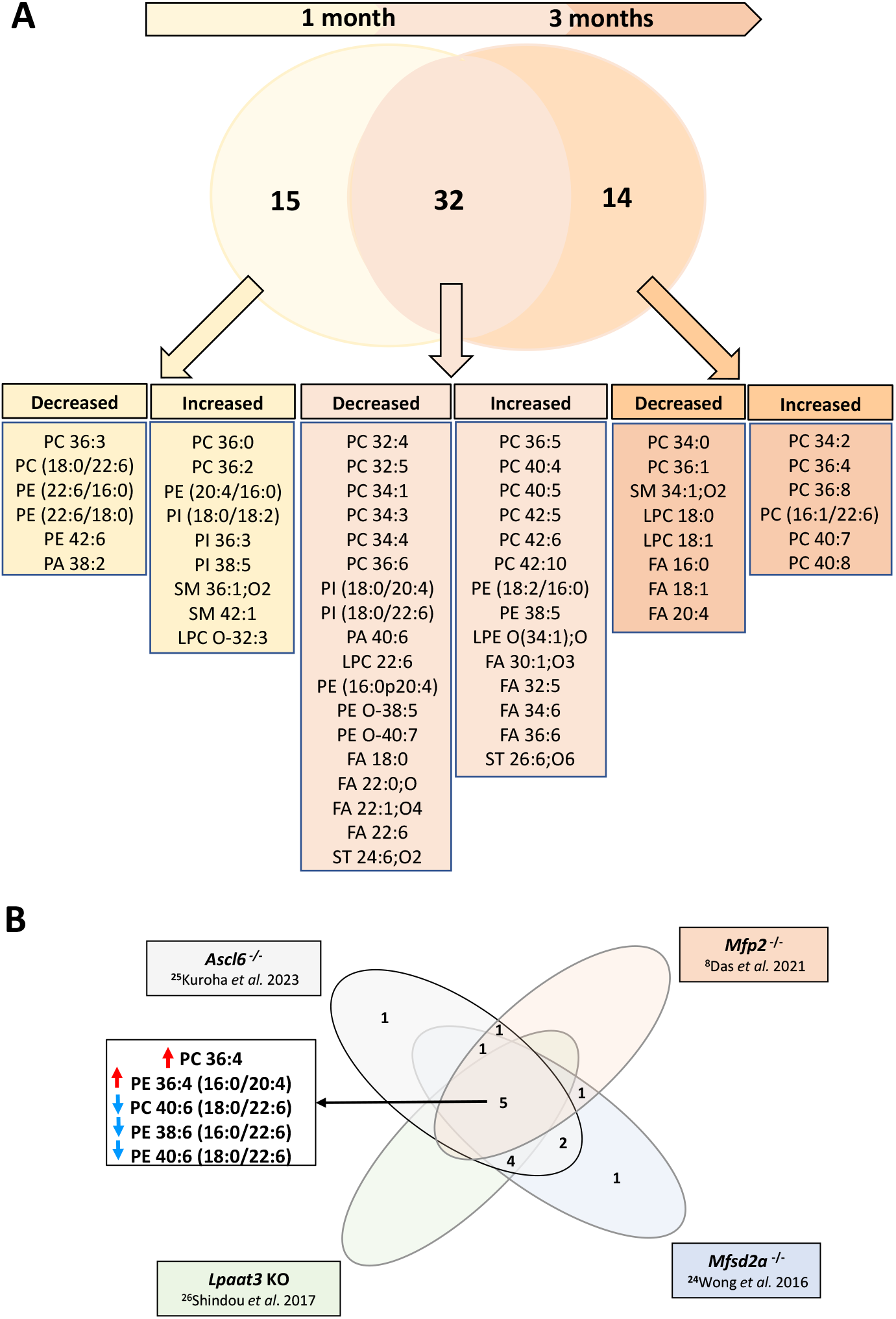
**A:** Venn diagram showing single lipid species significantly affected in PEX1-G844D mouse RPE at 1 and 3 months of age. **B:** Venn diagram comparing the overlap of lipid species affected in the RPE of PEX1-G844D mice and common to other genetic models of retinal degeneration with lipid dysregulation.

#### Identifying an RPE lipid signature that could be used as an early biomarker for RD

To understand whether the lipid changes observed in the RPE of PEX1-G844D mice could be directly linked to retinopathy progression, we investigated whether the lipids we identified by IMS in our model had already been described in different mouse models of RD. We identified from the scientific literature four genetically modified mouse models with altered lipids and RD: *Ascl6*^*-/-*^ [25], *Lpaat3* KO [26], *Mfp2*^*-/-*^ [8], and *Mfsd2a*^*-/-*^ [24]. Lipidomic analyses from these models showed 16 dysregulated lipids in common with those we identified using IMS in PEX1-G844D RPE (**Figure 5B**). Notably, we observed the same increase in PC 34:2 and PE 38:5, and the same decrease in DHA-associated lipids, PC 18:0/22:6 and PE 18:0/22:6, suggesting that these alterations are associated with retinopathy, and may be present more generally in RD.

## Discussion

In this study, we explored RPE structure, lipid dysregulation, and its implications in photoreceptor dysfunction in the PEX1-G844D mouse model of mild Zellweger spectrum disorder (ZSD). Our results reveal significant alterations in RPE lipid composition at early stages, which precede the onset of retinal changes. The RPE plays a crucial role in maintaining retinal homeostasis, and the lipid changes observed in this study highlight its importance as a key actor in understanding retinal dysfunction. Histological analysis of RPE flat mounts at 1 month of age revealed normal integrity of the PEX1-G844D RPE layer, with no significant lesions compared with control mice. However, lipid profiling by IMS identified over 40 lipids with significant differences in relative lipid densities, indicating changes in the membrane composition of RPE tissue in PEX1-G844D mice. These lipid alterations, notably the increase in VLCFAs and the decrease in plasmalogens and DHA-associated lipids, suggest that alterations in lipid metabolism may contribute to the development of photoreceptor dysfunction. Our results are consistent with known lipid abnormalities in ZSD patients, where elevated VLCFA levels and decreased plasmalogens have been observed [3].

Furthermore, the reduction of DHA-associated lipids is particularly relevant, as DHA is essential for neuronal development and the maintenance of photoreceptor integrity and function, and is protective against oxidative stress [27, 28]. Finally, DHA deficiency is a major contributor to RD in a mouse model of peroxisome β-oxidation deficiency [29]. The reduction in DHA-containing lipids in our PEX1-G844D model suggests an initial dysfunction associated with DHA deficiency, which may contribute to the observed photoreceptor dysfunction and possibly promote retinal damage.

This hypothesis was fully supported by our histological study of retinal tissue including RPE at 3 and 6 months of age revealing profound structural changes, with greatly enlarged and disorganized RPE cells, starting mainly in the dorsal pole and extending to the entire tissue over time. These structural abnormalities were accompanied by increased infiltration of inflammatory cells over time, indicating an active inflammatory response. In parallel, the observations of neural retina flat mounts and retinal cross-sections revealed decreased density of cone outer segments and outer nuclear thickness in general at 6 months, highlighting the consequences of an acute retinal stress in this specific area of the PEX1-G844D retina. Our data present for the first time in a ZSD mouse model a retinopathy phenotype dramatically affecting both the RPE and photoreceptor cells, with an inflammatory component. These results are consistent with the retinopathy phenotypes described in other models of RD associated with lipid dysregulation [8, 24-26].

The observed lipid accumulation in the dorsal pole could indicate region-specific stress or metabolic alterations, even if the cause or exact functional differences are not fully understood. This can help identify areas of the tissue that are more susceptible to changes and might guide further research into regional variations within the RPE. Possible explanation for the lipid imbalance in the dorsal pole of the RPE may be the higher metabolic activity of M cones in this region. M cones, which are more abundant in the dorsal retina [30-32], have increased energy requirements [33]. This increased metabolic activity leads to high lipid consumption, which can result in imbalance. Furthermore, the PEX1-G844D mutation can exacerbate this situation by impairing peroxisomal function, further disrupting lipid metabolism and homeostasis in the RPE. Therefore, the combination of increased lipid demand from metabolically active M cones and compromised peroxisomal function may contribute to the lipid imbalance observed in the dorsal pole of the RPE in our PEX1-G844D mice.

To understand whether the lipid changes observed in the RPE of PEX1-G844D mice could be driving retinpathy progression, we compared our results with those of different mouse models of retinal degeneration (RD). A literature comparison identified four genetically modified mouse models (*Ascl*^*-/-*^, *Lpaat3* KO, *Mfp2*^*-/-*^ and *Mfsd2a*^-/-^ with altered lipids and RD [8, 24-26]. Among these models, lipidomic analysis on retinal tissue revealed for each of them between 8 and 14 lipids in common with the PEX1-G844D RPE IMS data, in particular the increase in PC 34:2 and PE 38:5, and the decrease in DHA-associated lipids observed in all models. It should be noted that this common list is underestimated insofar as not all lipids identified in our model were analyzed in each of the studies. These similarities underline that the lipid alterations observed in our model may similarly play a role in retinopathy progression.

In this context, our results suggest that lipid dysregulation at early stages may contribute to the inflammation and retinal damage observed over time. Lipid profiling by IMS at 3 months of age revealed further changes in lipid composition, a persistent decrease in DHA and its associated lipids, and an exacerbated increase in VLCFAs. In addition, 14 new lipids showed further alterations, some of them highly accumulated in the dorsal pole, suggesting that their accumulation results from an inflammatory environment. While these lipid changes are associated with PEX1-G844D mutation-induced retinopathy, it is difficult to know whether they are related to the onset or progression of these disorders, or whether they are consequential to disease pathology. The lipid changes identified when tissues are damaged would, in turn, present a lipid signature of PEX1-G844D-induced retinopathy and be candidate biomarkers.

In addition to lipid dysregulation, we observed the formation of multiple vesicles or blebs accompanying the pronounced RPE actin cytoskeleton remodeling. Extracellular vesicles (EVs), which facilitate intercellular communication by transferring lipids, genetic material and proteins, may contribute to the propagation of lipid imbalances and inflammatory signals, exacerbating retinal degeneration [18]. Blebs, indicative of cellular stress, reflect the cellular response to lipid imbalance and structural changes in the RPE [34]. Identification of RPE-derived EVs and blebs in blood or plasma could serve as non-invasive biomarkers for monitoring disease progression and therapeutic response.

Our results highlight the importance of RPE in understanding PEX1-G844D-induced retinal dysfunction. Early lipid alterations and the subsequent inflammatory and structural defects suggest that RPE plays a key role in the progression of retinopathy. These findings pave the way for future research aimed at identifying which lipids drive the RD and could be novel therapeutic targets to intervene in the early stages of the disease. Although we focus here on lipid profiles, peroxisome functions are broad and include cellular redox maintenance and immune modulation [35, 36]. Deficiencies in these functions and their contribution to the RD phenotype warrants further investigation.

## Acknowledgements

We thank Frederic Fournelle for his MALDI method development efforts using murine retinal cryosections. This work was supported by a Canadian Institutes of Health Research (CIHR) Project Grant to NB, CA, and PC [201909PGT-427301-G-CFAC], and a Richard and Edith Strauss Foundation award to NB and CA

