## Supplemental table S1 for "Geographic characterization of RPE structure and lipid changes in the PEX1-p.Gly844Asp mouse model for Zellweger spectrum disorder"

| Lipid species (molecular species) | m/z | Control 1M | *PEX1-G844D* 1M | *P* value |
|---|---|---|---|---|
| PC 32:3 | 750,5068 | 48,49 ± 7,82 | 41,30 ± 1,95 | 0,4 |
| PC 32:4 | 748,4888 | 22,58 ± 2,23 | 15,27 ± 0,59 ** | 0,0053 |
| PC 32:5 | 746,4731 | 14,49 ± 0,51 | 9,83 ± 0,23 *** | 0,0001 |
| PC 34:0 | 762,6007 | 319,64 ± 9,35 | 305,51 ± 3,81 | 0,0725 |
| PC 34:1 | 760,5851 | 438,3 ± 30,38 | 370,09 ± 8,26 * | 0,0199 |
| PC 34:2 | 758,5694 | 517,12 ± 87,17 | 677,36 ± 74,58 | 0,0728 |
| PC 34:3 | 794,5097 | 30,27 ± 2,71 | 23,07 ± 0,92 * | 0,012 |
| PC 34:4 | 792,494 | 50,81 ± 1,83 | 38,69 ± 1,67 ** | 0,0011 |
| PC 36:0 | 812,614 | 158,33 ± 8,10 | 198,85 ± 11,97 ** | 0,0083 |
| PC 36:1 | 788,6164 | 211,54 ± 15,50 | 229,96 ± 13,78 | 0,1986 |
| PC 36:2 | 786,6007 | 267,7 ± 23,36 | 326,74 ± 19,32 * | 0,028 |
| PC 36:3 | 822,5492 | 32,92 ± 1,55 | 28,53 ± 0,80 * | 0,012 |
| PC 36:4 | 782,5694 | 794,85 ± 109,03 | 797,90 ± 46,03 | 0,9666 |
| PC 36:5 | 780,5538 | 303,76 ± 18,20 | 386,36 ± 20,34 ** | 0,0063 |
| PC 36:6 | 778,5381 | 79,73 ± 6,99 | 68,28 ± 0,60 * | 0,0475 |
| PC 36:8 | 774,5068 | 39,7 ± 3,36 | 35,95 ± 1,58 | 0,1556 |
| PC 38:4 | 810,6007 | 386,1 ± 30,12 | 387,79 ± 6,06 | 0,9288 |
| PC 38:6 | 806,5694 | 542,26 ± 89,43 | 410,63 ± 34,01 | 0,0758 |
| PC (16:1/22:6) | 804,5538 | 264,63 ± 66,06 | 315,89 ± 13,62 | 0,2585 |
| PC 40:4 | 860,614 | 6,08 ± 0,39 | 9,41 ± 0,56 ** | 0,0011 |
| PC 40:5 | 858,5983 | 13,99 ± 0,47 | 20,63 ± 0,63 *** | 0,0001 |
| PC (18:0/22:6) | 856,5827 | 43,75 ± 3,55 | 34,39 ± 2,89 * | 0,024 |
| PC 40:7 | 832,5851 | 25,12 ± 1,71 | 29,82 ± 3,06 | 0,081 |
| PC 40:8 | 830,5694 | 113,35 ± 22,68 | 127,71 ± 9,20 | 0,367 |
| PC 42:5 | 886,6296 | 2,34 ± 0,16 | 3,23 ± 0,28 ** | 0,0093 |
| PC 42:6 | 884,614 | 3,43 ± 0,19 | 4,13 ± 0,37 * | 0,0424 |
| PC 42:10 | 892,5253 | 1,75 ± 0,09 | 2,33 ± 0,22 * | 0,0127 |
| PE (18:2/16:0) | 714,5079 | 17,30 ± 36,29 | 38,81 ± 4,95 ** | 0,0019 |
| PE (20:4/16:0) | 738,5079 | 87,88 ± 36,29 | 164,12 ± 10,99 * | 0,0253 |
| PE 38:5 | 764,5236 | 35,47 ± 8,36 | 51,74 ± 1,74 * | 0,03 |
| PE (22:6/16:0) | 762,5079 | 92,1 ± 15,96 | 50,52 ± 10,83 * | 0,0202 |
| PE (22:6/18:0) | 790,5392 | 145,64 ± 31,16 | 68,45 ± 18,32 * | 0,0209 |
| PE 42:6 | 818,5705 | 17,8 ± 2,52 | 12,19 ± 1,30 * | 0,0268 |
| PI (18:0/18:2) | 861,5499 | 15,12 ± 1,18 | 26,16 ± 0,76 *** | 0,0002 |
| PI 36:3 | 859,5342 | 13,51 ± 2,22 | 20,73 ± 1,74 * | 0,0114 |
| PI (16:0/20:4) | 857,5186 | 42,65 ± 12,34 | 57,63 ± 3,40 | 0,1125 |
| PI (18:0/20:4) | 885,5499 | 411,37 ± 122,79 | 189,31 ± 19,69 * | 0,0365 |
| PI 38:5 | 883,5342 | 22,44 ± 4,41 | 30,99 ± 1,07 * | 0,0309 |
| PI (18:0/22:6) | 909,5499 | 23,32 ± 4,07 | 15,65 ± 1,99 * | 0,0426 |
| PA 38:2 | 763,492 | 36,49 ± 6,45 | 22,41 ± 4,28 * | 0,0345 |
| PA 40:6 | 747,497 | 33,27 ± 3,79 | 15,14 ± 1,49 ** | 0,0015 |
| SM 34:1;O2 | 725,5568 | 245,31 ± 43,03 | 214,97 ± 34,73 | 0,3957 |
| SM 36:1;O2 | 731,6061 | 19,03 ± 1,51 | 28,76 ± 1,67 ** | 0,0017 |
| SM 42:1 (d18:0/24:1) | 837,682 | 29,25 ± 2,16 | 38,15 ± 1,58 ** | 0,0045 |
| PG (22:6/22:6) | 865,5025 | 6,57 ± 2,27 | 3,53 ± 0,93 | 0,0981 |
| LPC 16:0 | 496,33 | 82,44 ± 19,33 | 76,95 ± 14,73 | 0,7155 |
| LPC 18:0 | 524,37 | 27,12 ± 9,62 | 18,93 ± 5,11 | 0,2629 |
| LPC 18:1 | 522,35 | 10,8 ± 1,70 | 7,64 ± 1,14 | 0,0553 |
| LPC 18:2 | 520,33 | 15,67 ± 3,17 | 16,90 ± 2,64 | 0,6315 |
| LPC 20:4 | 544,33 | 27,25 ± 7,46 | 19,73 ± 1,54 | 0,1621 |
| LPC 22:0 | 584,39 | 6,80 ± 1,13 | 8,28 ± 1,02 | 0,1677 |
| LPC 22:6 | 568,33 | 24,90 ± 3,05 | 12,40 ± 1,46 ** | 0,0031 |
| LPE O(34:0);O | 742,5523 | 54,34 ± 20 | 68,03 ± 10,96 | 0,3572 |
| LPE O(34:1);O | 740,5367 | 6,43 ± 1,39 | 11,80 ± 1,37 ** | 0,0089 |
| LPC O-32:3 | 698,54 | 40,78 ± 4,87 | 29,31 ± 3,26 * | 0,0275 |
| PE O-36:5 | 722,51 | 59,46 ± 8,22 | 29,13 ± 2,87 ** | 0,0038 |
| PE O-38:5 | 750,54 | 44,18 ± 4,12 | 19,14 ± 0,55 *** | 0,0005 |
| PE O-40:7 | 774,54 | 29,97 ± 1,58 | 14,33 ± 1,6 *** | 0,0003 |
| PA O-36:1 | 687,53 | 249,67 ± 10,85 | 219,19 ± 34,45 | 0,2177 |
| PA O-42:1 | 771,62 | 183,88 ± 20,68 | 142,5 ± 17,52 | 0,0573 |
| PA O-42:2 | 769,61 | 92,72 ± 19,94 | 60,64 ± 8,03 | 0,061 |
| FA 16:0 | 363,14 | 16,87 ± 3,78 | 16,08 ± 0,73 | 0,767 |
| FA 18:0 | 391,17 | 37,7 ± 4,7 | 25,63 + 0,73 * | 0,0118 |
| FA 18:1 | 389,16 | 42,47 ± 2,89 | 37,53 ± 3,86 | 0,151 |
| FA 18:2 | 387,14 | 14,41 ± 2,28 | 18,76 ± 2,13 | 0,0733 |
| FA 20:4 | 411,14 | 42,33 ± 3,35 | 34,89 ± 3,86 | 0,0655 |
| FA 22:0;O | 463,23 | 13,28 ± 1,62 | 3,83 ± 1,01 ** | 0,001 |
| FA 22:1;O4 | 509,2 | 18,84 ± 5,95 | 5,42 ± 0,55 * | 0,0177 |
| FA 22:6 | 435,14 | 79,92 ± 8,05 | 25,18 ± 7,51 *** | 0,001 |
| FA 30:1;O3 | 605,33 | 0,672 ± 0,06 | 7,382 ± 1,23 *** | 0,0007 |
| FA 32:5 | 577,31 | 0,51 ± 0,05 | 4,81 ± 1,08 ** | 0,0023 |
| FA 34:6 | 603,33 | 0,66 ± 0,1 | 3,97 ± 0,91 ** | 0,0033 |
| FA 36:6 | 631,36 | 0,52 ± 0,03 | 1,95 ± 0,4 ** | 0,0035 |
| ST 24:6;O2 | 457,12 | 14,07 ± 2,22 | 3,44 ± 1,85 ** | 0,0031 |
| ST 26:6;O6 | 549,14 | 0,51 ± 0,05 | 1,29 ± 0,23 ** | 0,0045 |
| ST 27:1;O | 493,25 | 13,61 ± 2,52 | 11 ± 0,68 | 0,1579 |

**Supplemental Table S1**: **Changes in the RPE lipid composition in 1-month-old PEX1-G844D mice.** Imaging Mass Spectrometric (IMS) analysis of the lipid composition in RPE tissue from 1-month-old WT and PEX1-G844D mice identified 76 lipid species, 47 of which showed a significant change in abundance (N=3, *p<0.05, **p<0.01, ***p<0.001).
