## Supplemental table S2 for "Geographic characterization of RPE structure and lipid changes in the PEX1-p.Gly844Asp mouse model for Zellweger spectrum disorder"

| Lipid species (molecular species) | m/z | Control 3M | *PEX1-G844D* 3M | *P* value |
|---|---|---|---|---|
| PC 32:3 | 750,5068 | 41,53 ± 0,33 | 38,64 ± 6,01 | 0,4523 |
| PC 32:4 | 748,4888 | 23,25 ± 3,52 | 16,81 ± 1,16 * | 0,0396 |
| PC 32:5 | 746,4731 | 15,8 ± 2,10 | 11 ± 1,30 * | 0,0281 |
| PC 34:0 | 762,6007 | 296,45 ± 27,39 | 216,3 ± 34,61 * | 0,0347 |
| PC 34:1 | 760,5851 | 419,07 ± 15,41 | 289,34 ± 28,90 ** | 0,0024 |
| PC 34:2 | 758,5694 | 273,48 ± 65,09 | 608,88 ± 103,63 ** | 0,009 |
| PC 34:3 | 794,5097 | 32,6 ± 1,93 | 21,69 ± 1,23 ** | 0,0012 |
| PC 34:4 | 792,494 | 46,57 ± 1,55 | 32,94 ± 3,47 ** | 0,0034 |
| PC 36:0 | 812,614 | 135,59 ± 11,18 | 164,59 ± 16,97 | 0,0688 |
| PC 36:1 | 788,6164 | 229,19 ± 17,36 | 158,16 ± 18,73 ** | 0,0085 |
| PC 36:2 | 786,6007 | 193,71 ± 37,96 | 270,53 ± 32,11 | 0,0555 |
| PC 36:3 | 822,5492 | 26,33 ± 1,33 | 30,69 ± 3,60 | 0,12 |
| PC 36:4 | 782,5694 | 745,17 ± 32,95 | 947,79 ± 84,25 * | 0,0178 |
| PC 36:5 | 780,5538 | 212,85 ± 24,49 | 341,85 ± 59,68 * | 0,0257 |
| PC 36:6 | 778,5381 | 92,18 ± 8,6 | 68,28 ± 7,89 * | 0,0239 |
| PC 36:8 | 774,5068 | 30,87 ± 0,83 | 44,25 ± 5,10 * | 0,0109 |
| PC 38:4 | 810,6007 | 355,41 ± 25,17 | 368,14 ± 56,88 | 0,741 |
| PC 38:6 | 806,5694 | 354,32 ± 80,32 | 425,29 ± 52,22 | 0,2687 |
| PC (16:1/22:6) | 804,5538 | 168,99 ± 19,53 | 340,4 ± 27,27 *** | 0,0009 |
| PC 40:4 | 860,614 | 7,78 ± 0,66 | 11,72 ± 0,14 *** | 0,0006 |
| PC 40:5 | 858,5983 | 14,7 ± 1,81 | 24,99 ± 2,21 ** | 0,0033 |
| PC (18:0/22:6) | 856,5827 | 46,6 ± 6,84 | 44,98 ± 8,40 | 0,8079 |
| PC 40:7 | 832,5851 | 115,02 ± 14,75 | 166,63 ± 25,29 * | 0,0379 |
| PC 40:8 | 830,5694 | 69,4 ± 19,66 | 119,32 ± 11,45 * | 0,0191 |
| PC 42:5 | 886,6296 | 2,97 ± 0,39 | 3,96 ± 0,32 * | 0,0272 |
| PC 42:6 | 884,614 | 4,64 ± 0,28 | 5,65 ± 0,38 * | 0,0211 |
| PC 42:10 | 892,5253 | 2,52 ± 0,32 | 3,6 ± 0,47 * | 0,0295 |
| PE (18:2/16:0) | 714,5079 | 13,06 ± 3,05 | 26,18 ± 3,82 ** | 0,0097 |
| PE (20:4/16:0) | 738,5079 | 114,47 ± 28,80 | 147,83 ± 20,45 | 0,1772 |
| PE 38:5 | 764,5236 | 38,73 ± 7,36 | 64,01 ± 8,32 * | 0,0169 |
| PE (22:6/16:0) | 762,5079 | 145,35 ± 38,74 | 130,32 ± 6,07 | 0,5429 |
| PE (22:6/18:0) | 790,5392 | 179,49 ± 63,29 | 212,74 ± 27,80 | 0,4517 |
| PE 42:6 | 818,5705 | 17,37 ± 5,87 | 18,73 ± 2,12 | 0,7261 |
| PI (18:0/18:2) | 861,5499 | 15,27 ± 3,44 | 22,54 ± 3,13 | 0,0537 |
| PI 36:3 | 859,5342 | 15,99 ± 2,49 | 20,24 ± 3,90 | 0,1871 |
| PI (16:0/20:4) | 857,5186 | 67,61 ± 19,26 | 72,97 ± 12,25 | 0,7047 |
| PI (18:0/20:4) | 885,5499 | 493,07 ± 107,84 | 290,64 ± 21,33 * | 0,0332 |
| PI 38:5 | 883,5342 | 18,69 ± 3,05 | 23,2 ± 2,58 | 0,1222 |
| PI (18:0/22:6) | 909,5499 | 37,9 ± 6,88 | 21,61 ± 4,56 * | 0,0268 |
| PA 38:2 | 763,492 | 58,52 ± 15,84 | 53,14 ± 1,67 | 0,5897 |
| PA 40:6 | 747,497 | 47,75 ± 10,05 | 21,09 ± 1,12 * | 0,0103 |
| SM 34:1;O2 | 725,5568 | 260,94 ± 53,93 | 123,99 ± 12,21 * | 0,0127 |
| SM 36:1;O2 | 731,6061 | 16,39 ± 1,90 | 19,01 ± 3,67 | 0,3337 |
| SM 42:1 (d18:0/24:1) | 837,682 | 55,96 ± 2,54 | 55,67 ± 2,25 | 0,8891 |
| PG 44:12 (22:6/22:6) | 865,5025 | 3,57 ± 0,43 | 4,35 ± 0,35 | 0,0718 |
| LPC 16:0 | 496,33 | 84,36 ± 13,96 | 72,22 ± 1,54 | 0,2089 |
| LPC 18:0 | 524,37 | 57,8 ± 20,63 | 19,77 ± 2,15 * | 0,0337 |
| LPC 18:1 | 522,35 | 21,75 ± 2,34 | 6,56 ± 1,23 *** | 0,0006 |
| LPC 18:2 | 520,33 | 15,86 ± 8,17 | 10,68 ± 5,69 | 0,4188 |
| LPC 20:4 | 544,33 | 17,93 ± 8,05 | 10,24 ± 3,04 | 0,1963 |
| LPC 22:0 | 584,39 | 4,11 ± 0,26 | 4,36 ± 0,85 | 0,6576 |
| LPC 22:6 | 568,33 | 23,02 ± 4,92 | 9,28 ± 1,05 ** | 0,0091 |
| LPE O(34:0);O | 742,5523 | 87,46 ± 27,54 | 119,17 ± 23,27 | 0,2023 |
| LPE O(34:1);O | 740,5367 | 12,22 ± 2,65 | 19,43 ± 3,21 * | 0,04 |
| LPC O-32:3 | 698,54 | 21,49 ± 3,42 | 16,76 ± 2,46 | 0,1237 |
| PE O-36:5 | 722,51 | 60,62 ± 8,02 | 28,49 ± 5,5 ** | 0,0046 |
| PE O-38:5 | 750,54 | 56,66 ± 3,59 | 25,56 ± 5,41 ** | 0,0011 |
| PE O-40:7 | 774,54 | 46,08 ± 14,94 | 21,38 ± 1,77 * | 0,0467 |
| PA O-36:1 | 687,53 | 147,56 ± 29,08 | 106,39 ± 8,38 | 0,078 |
| PA O-42:1 | 771,62 | 96,11 ± 40,94 | 50,61 ± 8,78 | 0,1329 |
| PA O-42:2 | 769,61 | 54,28 ± 21,55 | 34,88 ± 5,19 | 0,2042 |
| FA 16:0 | 363,14 | 22,39 ± 3,13 | 14 ± 2,25 * | 0,0195 |
| FA 18:0 | 391,17 | 44,87 ± 11,20 | 20,62 ± 3,07 * | 0,0224 |
| FA 18:1 | 389,16 | 50,03 ± 8,85 | 29,59 ± 6,11 * | 0,0301 |
| FA 18:2 | 387,14 | 17,59 ± 1,88 | 15,99 ± 3,89 | 0,5555 |
| FA 20:4 | 411,14 | 44,25 ± 3,70 | 24,92 ± 5,85 ** | 0,0084 |
| FA 22:0;O | 463,23 | 8,48 ± 0,83 | 3,81 ± 0,53 ** | 0,0012 |
| FA 22:1;O4 | 509,2 | 23,32 ± 2,85 | 4,1 ± 0,36 *** | 0,0003 |
| FA 22:6 | 435,14 | 99,85 ± 20,62 | 12,52 ± 1,15 ** | 0,0018 |
| FA 30:1;O3 | 605,33 | 0,89 ± 0,08 | 30,91 ± 7,59 ** | 0,0024 |
| FA 32:5 | 577,31 | 0,73 ± 0,08 | 14,57 ± 3,77 ** | 0,0032 |
| FA 34:6 | 603,33 | 0,87 ± 0,14 | 15,28 ± 2,29 *** | 0,0004 |
| FA 36:6 | 631,36 | 0,6 ± 0,05 | 5,28 ± 1,80 * | 0,0107 |
| ST 24:6;O2 | 457,12 | 12,46 ± 3,05 | 1,57 ± 0,17 ** | 0,0035 |
| ST 26:6;O6 | 549,14 | 0,72 ± 0,10 | 2,16 ± 0,43 ** | 0,0048 |
| ST 27:1;O | 493,25 | 214,03 ± 48,56 | 166,93 ± 18,16 | 0,1907 |

**Supplemental Table S2: Changes in the RPE lipid composition in 3-month-old PEX1-G844D mice.** Imaging mass spectrometric (IMS) analysis of the lipid composition of RPE tissue from 3-month-old WT and PEX1-G844D mice identified 72 lipid species, 46 of which showed a significant change in abundance (N=3, *p<0.05, **p<0.01, ***p<0.001).
